# A Structural Model of Transcriptional Activation by Mediator

**DOI:** 10.64898/2026.10.01.756097

**Authors:** Andrey Feklistov, Roger D. Kornberg

## Abstract

Mediator integrates regulatory signals from transcription factors (TFs) and controls RNA polymerase II (Pol II) activity in eukaryotes. Its Tail module serves as the primary hub for TF activation domains (ADs), but the mechanistic logic coupling this distal engagement to transcriptional output is unclear. Here we model an extensive set of human AD–Tail interactions with AlphaFold 3 and infer how they may influence the architecture of the Mediator–Pol II complex. Across the ensemble, ADs recurrently engage a restricted set of surfaces spanning multiple Tail subunits, revealing several previously unrecognized sites. The ensemble map recovers established activator-binding regions and is consistent with prior biochemical and structural data. Our modeling suggests how multivalent AD binding modulates the conformational landscape of the Tail to favor an initiation-competent Pol II state, providing a structural framework for tunable gene activation through Mediator.

## Introduction

The defining mechanistic problem of transcriptional activation is how DNA-bound activators alter the probability of transcription initiation at a promoter. Activators enlist a diverse repertoire of co-regulators, but Mediator stands out for its broad ability to integrate their inputs into quantitative changes in RNA output: shaping burst frequency, inducible dynamic range and synergistic responses to multiple activators [Malik and Roeder, 2023; Zhou et al., 2025]. This regulatory capacity stems from the privileged position of Mediator within the preinitiation complex (PIC): it engages RNA polymerase II (Pol II) directly and extensively along the back surface of the enzyme. Its Head and Middle modules contact the clamp and neighboring regions whose motions shape the Pol II active-site cleft, whereas the Tail projects from this Pol II-engaged core and serves as the principal interaction platform for transcription-factor (TF) activation domains (ADs) [Abdella et al., 2021; Rengachari et al., 2021]. This architecture poses a long-range signaling challenge: weak and transient AD contacts are registered by the flexible Tail, but their consequences must be transmitted across Mediator to the Pol II initiation machinery.

ADs are generally intrinsically disordered and enriched in acidic and bulky hydrophobic residues, but lack a single conserved interaction motif. High-throughput tiling and mutational analyses have mapped hundreds of human ADs and identified short sequence segments that, when deleted, can either reduce or enhance activation [DelRosso et al., 2023]. Biophysical studies further show that AD–Mediator interactions are weak, dynamic and structurally heterogeneous, supporting a multivalent model in which distributed AD elements contribute to binding [Brzovic et al., 2011; Tuttle et al., 2018; Sanborn et al., 2021].

Cryo-EM analysis of human Mediator resolved the Tail in two markedly different conformations: Extended and Bent [Chen et al., 2021]. We previously proposed that the Extended architecture is associated with a Pol II cleft geometry compatible with promoter DNA melting and transcription initiation [Kornberg and Feklistov, 2025]. This model envisions a spatial correspondence between interaction elements within ADs and binding surfaces across the Tail that could bias its conformation and affect the geometry of the Mediator–Pol II interface.

Mapping how diverse ADs engage the human Mediator Tail and how these interactions relate to its conformational states is not currently tractable by experimental structural methods. We therefore used AlphaFold 3 (AF3) [Abramson et al., 2024] to examine these interactions at scale, modeling the human Mediator Tail with nearly 300 functionally mapped human AD constructs [DelRosso et al., 2023]. The predicted AD–Tail interfaces generally carried low confidence and did not converge on a unique AD orientation or residue register. Accordingly, we focused on recurrent patterns of contact with Tail surfaces across the ensemble, without assigning structural significance to individual poses [Omidi et al., 2024]. We then examined this surface map in light of systematic deletion data [DelRosso et al., 2023] and the experimentally determined Extended and Bent Tail structures [Chen et al., 2021].

The resulting ensemble reveals that ADs repeatedly engage a restricted set of surfaces spanning multiple Tail subunits. These include established activator-binding regions on MED15 and MED23 together with previously unrecognized recurrent receptor sites on MED16, MED24 and the MED25-containing tunnel. Systematic deletion analysis reinforces this organization: AD segments that frequently contact Mediator are often required for activation, whereas reducing the sequence spacing between such segments can enhance it. Together, these observations support a model in which multivalent engagement across the Tail favors the Extended conformation. We further suggest how this conformational bias may be transmitted through Mediator to influence Pol II cleft geometry and the probability of productive initiation.

## Results

### ADs repeatedly engage multiple sites on one face of the Mediator Tail

Structures of Mediator that include the Tail module have been determined in budding yeast, a thermophilic fungus and mammals [Gorbea Colón et al., 2023; Zhang et al., 2021; Chen et al., 2021]. Among these systems, human Mediator was particularly well suited to our analysis, with a major Tail rearrangement captured by cryo-EM [Chen et al., 2021] and a large repertoire of activation domains mapped by systematic screens [DelRosso et al., 2023]. MED15 organization further favored the human complex. In budding yeast, multiple activator-binding domains (ABDs) are distributed through a long disordered region, whereas one structured ABD of human MED15 lies in its central region, close to the resolved Tail core [Zhao et al., 2013; Yu et al., 2025; Chen et al., 2021]. This compact arrangement also defined our modeling boundary: we retained the Tail-proximal structured region of MED15 and excluded the more distal low-complexity sequence. We then modeled AD engagement using a seven-chain Tail receptor containing this MED15 region, the MED27 and MED29 segments forming the shared helical bundle, MED16, MED24 and MED23, and the N-terminal VWA domain of MED25. The long unresolved MED25 region containing the canonical ACID/VBD was not modeled.

Across the 293 AF3 prediction sets, Tail geometry closely matched the Extended-state Tail resolved by cryo-EM (median Cα RMSD = 1.89 Å; IQR, 1.79–1.96 Å; 3,328 paired Cα atoms). AF3 did not recover the Bent state even with templates derived from its cryo-EM structure. Extended-like Tail geometries are represented in many more mammalian structures than Bent, an imbalance that may contribute to this AF3 preference. We therefore analyze AD contacts in the Extended-state geometry without using AF3 state preference to infer an effect of AD binding on the Tail conformational equilibrium.

We modeled 293 human AD constructs from the functionally mapped repertoire [DelRosso et al., 2023], generating five AF3 models per construct (1,465 models total). Models in which AF3 used the AD to complete Mediator structure missing from the receptor construct (a structural-inpainting artifact), or threaded the AD through sterically implausible openings, were excluded; affected local contacts were masked when the remainder of a model was usable. This left 1,139 models from 234 AD constructs for analysis. After aligning the AF3 models on the Tail core, AD positions revealed a pronounced spatial asymmetry. Although exposed hydrophobic patches occur across much of the Tail surface, modeled AD traffic was largely confined to one face, with little occupancy on the opposite surface (Fig. 1a). The distribution therefore did not simply follow exposed hydrophobic surface. AD traffic on this face converged at discrete sites on MED15, MED16, MED23, MED24 and MED25. Despite substantial variation in AD orientation and residue register, the same Tail surfaces were repeatedly sampled across the ensemble, supporting analysis at the level of receptor-site engagement rather than individual docked poses.

**Figure 1.**
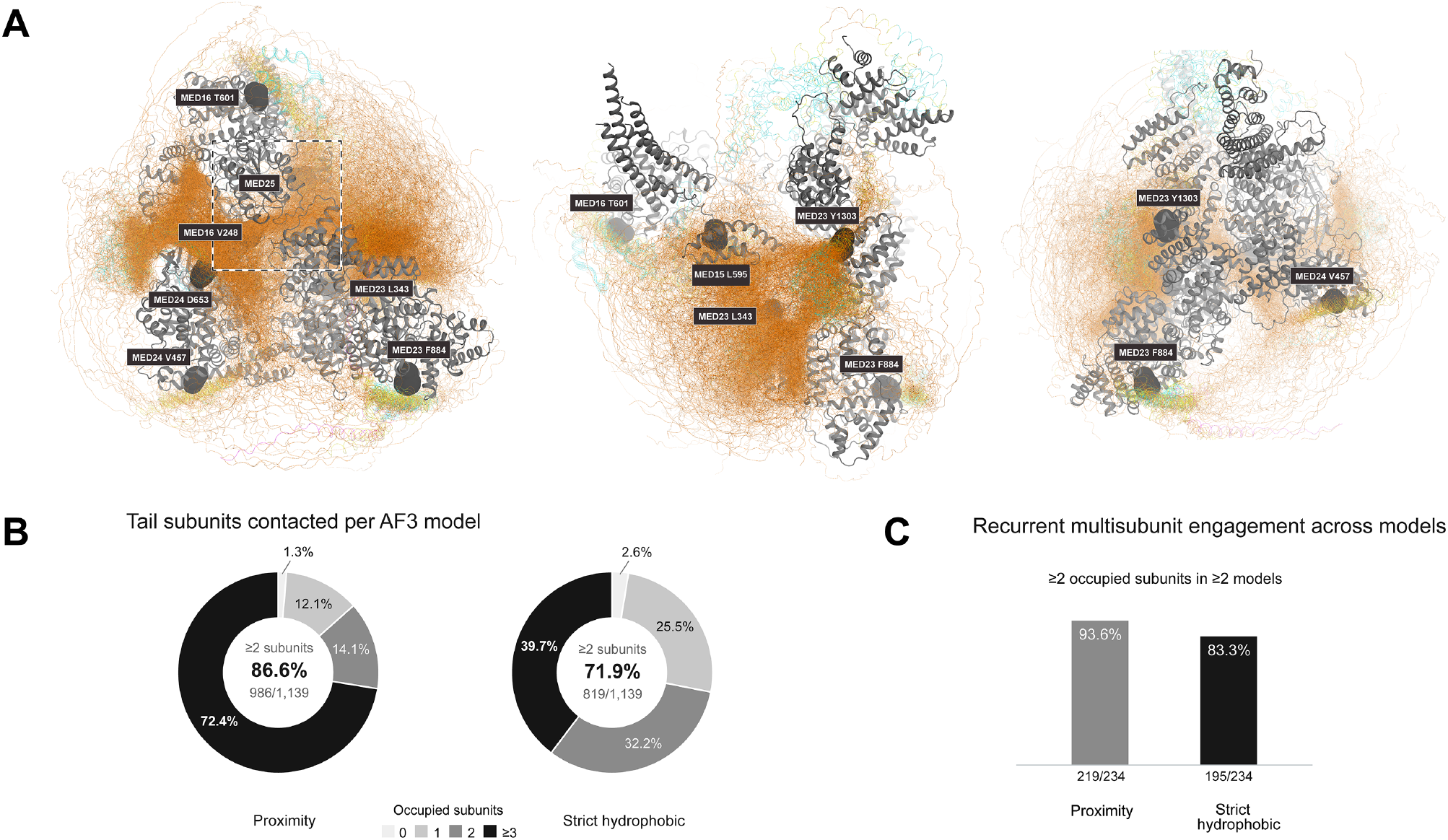
Activation domains engage multiple Mediator Tail subunits and converge on recurrent receptor sites. **(A) Three-dimensional map of AD engagement across the Mediator Tail.** Three views of the aligned AF3 ensemble are shown. The Mediator Tail is dark gray, and ADs are colored by AF3 pLDDT: orange, <50; yellow, 50–70; cyan, 70–90; and blue, >90. Labeled spheres indicate recurrent receptor sites on MED15 (L595), MED16 (V248, T601), MED24 (V457, D653) and MED23 (L343, F884, Y1303). The dashed box highlights the MED16–MED25 tunnel, a major recurrent receptor site between the MED16 β-propeller rim and MED25 VWA domain, centered near MED16 V248. The canonical MED25 ACID/VBD was not modeled. **(B) Tail subunits contacted per AF3 model.** Distribution of the number of distinct Tail subunits contacted across 1,139 usable AF3 models. Donut sectors, from light gray to black, indicate contact with 0, 1, 2 or ≥3 subunits. At least two subunits were contacted by 986 models (86.6%) by proximity and 819 models (71.9%) by the strict hydrophobic criterion. **(C) Recurrent multisubunit engagement across models.** Recurrence was evaluated across 234 AD constructs, each represented by two to five usable models. A construct was classified as recurrent when at least two models each contacted at least two Tail subunits. This criterion was met by 219 constructs (93.6%) by proximity and 195 constructs (83.3%) by the strict hydrophobic criterion. Proximity contacts were defined by Cα–Cα distances <12 Å; strict hydrophobic contacts by W/F/Y/L/I/V/M side-chain distances ≤4.5 Å.

Major sites of AD traffic recovered experimentally established activator-binding regions on MED15 and MED23. The agreement is especially direct for MED23, where L343 marks the peak of AD traffic and lies within the TF Elk-1–MED23 interface resolved by cryo-EM [Monté et al., 2025]. Beyond these established regions, the ensemble identified several previously unrecognized recurrent sites. A major receptor site occupied a tunnel formed between the MED16 β-propeller rim and the MED25 VWA domain, centered near V248. Additional traffic mapped to a separate helical site on MED16 centered around T601, to sites on MED24 near V457 and D653, and to distal MED23 sites near F884 and Y1303. MED16 itself has independent precedent as an activator-contacting subunit: the Gcn4 N-terminal AD crosslinks to yeast Med16, and human MED16 binds NRF2 directly [Herbig et al., 2010; Sekine et al., 2016], although neither study mapped a discrete binding site on MED16. Because the canonical MED25 ACID/VBD was not included in our model, the MED16–MED25 tunnel represents a distinct receptor architecture from the well-characterized MED25 activator-binding domain.

AD contacts within individual models frequently spanned more than one Tail subunit. We quantified this pattern at two levels: within individual AF3 models and among models for each AD construct. First, we asked how many Tail subunits were contacted within each of the 1,139 usable models. By a proximity criterion (Cα–Cα <12 Å), 986 models (86.6%) contacted at least two subunits; 819 (71.9%) did so under the stricter side-chain hydrophobic criterion (W/F/Y/L/I/V/M side-chain distances ≤4.5 Å). We next asked whether this pattern recurred among the models generated for the same AD construct. An AD construct was scored as recurrent when at least two of its usable models each contacted two or more Tail subunits; 219 of 234 constructs (93.6%) met this criterion by proximity and 195 (83.3%) by the strict criterion (Fig. 1b,c). Thus, multisubunit engagement was not confined to isolated models or a small subset of ADs, but was repeatedly recovered for most AD constructs.

### Deletion phenotypes reveal interaction elements and linkers within ADs

We next asked whether the structural patterns emerging from the AF3 ensemble were reflected in independent functional measurements of the underlying AD sequences. We mapped the systematic deletion data of DelRosso et al. onto the modeled constructs, distinguishing activation-impairing deletions, including those that abolished activity, from neutral and activation-enhancing deletions [DelRosso et al., 2023]. In the spatial projections, regions required for activation clustered around high-traffic regions of the aggregate AF3 contact map (Fig. 2a). Activation-enhancing deletions appeared less centered on these regions (Fig. 2b), prompting a quantitative comparison in AD sequence space.

**Figure 2.**
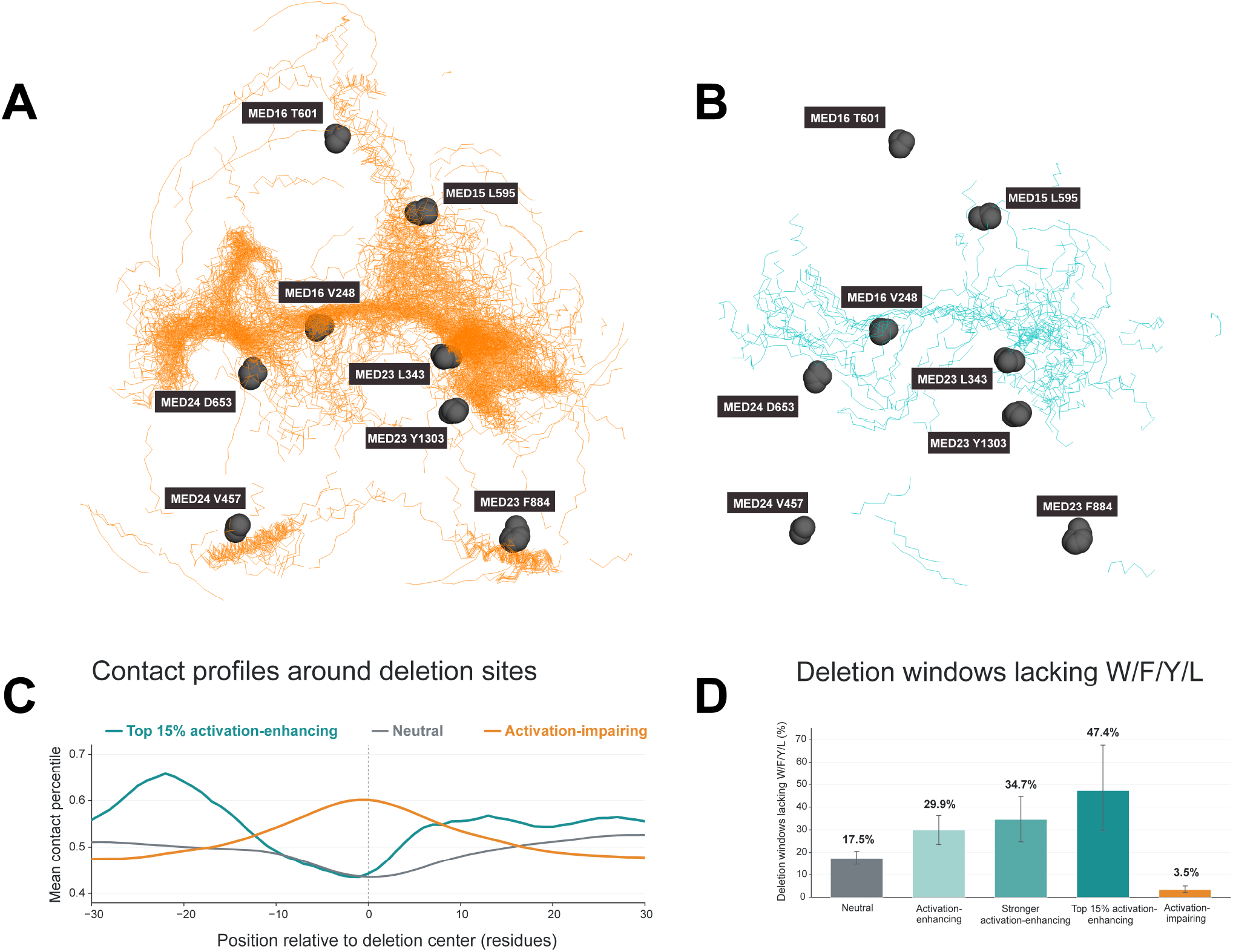
Activation-impairing and activation-enhancing deletions occupy distinct AD contact environments. (A) AD regions required for activation in the AF3 ensemble. Orange traces show these regions in the aligned AF3 models; the Mediator Tail is not shown; spheres mark recurrent Tail receptor sites. (B) AD regions removed by activation-enhancing deletions in the AF3 ensemble. Teal traces show these regions in the aligned AF3 models; the Mediator Tail is not shown; spheres mark recurrent Tail receptor sites. (C) **Contact profiles around deletion sites.** Per-residue Mediator contact was ranked within each AD, profiles were aligned at deletion centers, and mean contact percentile was plotted for activation-impairing, neutral and top 15% activation-enhancing deletions. Activation-impairing deletions center on a local contact maximum, whereas the top 15% activation-enhancing deletions center on lower-contact sequence flanked by higher-contact regions. (D) **Deletion windows lacking W/F/Y/L.** Fraction of neutral, activation-enhancing, stronger activation-enhancing, top 15% activation-enhancing and activation-impairing deletion windows lacking W/F/Y/L residues. Whiskers indicate 95% confidence intervals. Class definitions are given in Methods.

For this comparison, we aligned contact profiles on each deletion center and compared relative Mediator contact across the surrounding sequence. Activation-impairing deletions were centered on a pronounced local maximum of Mediator contact, consistent with direct removal of sequence involved in the interaction (Fig. 2c). The top 15% of activation-enhancing deletions showed the opposite profile: Mediator contact was lower at their centers and increased in the flanking sequence. The centers of this subset were similar to those of neutral deletions, indicating that low contact at the deletion center is not unique to the activation-enhancing class. The top 15% of activation-enhancing deletions therefore tend to remove relatively low-contact sequence positioned between or beside higher-contact interaction elements.

To further characterize the low-contact sequence targeted by activation-enhancing deletions, we examined the intervals between local peaks of Mediator contact. Measured along the modeled AD constructs, these deletions occurred within longer interpeak intervals than the reference distribution: intervals overlapped by activation-enhancing deletions had a median length of 43 residues, compared with 28 residues across all interpeak intervals and 47 residues for the stronger activation-enhancing subset (Extended Data Fig. 1). Sequence composition revealed a second difference. Activation-enhancing deletion windows were more often devoid of W/F/Y/L residues than neutral deletions, whereas activation-impairing deletions rarely lacked these residues; the distinction became stronger when the comparison was restricted to interpeak sequence (Fig. 2d). W/F/Y/L residues are strongly enriched in functionally required AD sequence [DelRosso et al., 2023], and those regions also show preferential Mediator contact in the AF3 ensemble. Together, these observations suggest that activation-enhancing deletions often shorten longer linkers separating interaction elements, whereas activation-impairing deletions preferentially remove the interaction elements themselves.

These results provide a structural context for the observation that many human ADs contain several short regions required for activation: multiple functionally important interaction elements can be distributed along the same disordered chain [DelRosso et al., 2023; Tuttle et al., 2018; Sanborn et al., 2021]. Consistent with this organization, residues with high Mediator contact were enriched in regions of higher local AF3 confidence (pLDDT), and activation-impairing deletions preferentially removed these better-defined elements (Extended Data Fig. 2). By contrast, activation-enhancing deletions showed no reproducible depletion of high-pLDDT or helical sequence relative to neutral deletions (Source Data for Extended Data Fig. 2). Shortening sequence between or beside the interaction elements could alter their spacing or effective concentration, reduce the conformational search required for multivalent engagement, or remove sequence that otherwise limits productive contact; the present data do not distinguish among these possibilities. Together with the multisubunit occupancy described above, the deletion phenotypes are consistent with activation depending on the relative organization of multiple interaction elements within an AD.

### AD traffic maps onto a Tail surface remodeled between Mediator states

Across the AF3 ensemble, individual ADs commonly contacted multiple Tail subunits, and this pattern recurred across most constructs (Fig. 1b,c). This raised a more specific question: were these contacts distributed broadly across the Tail, or did certain receptor sites tend to be engaged together? To examine these relationships, we mapped the recurrent Tail receptor sites onto the Extended Tail structure (Fig. 3a) and represented co-engagement as a weighted network (Fig. 3b). Node area reflects AD traffic at each site, and edge width the number of AF3 models in which the same AD contacted both sites under the strict hydrophobic criterion. The network was strongly non-uniform. The most frequent connection linked MED16 V248, at the center of the MED16–MED25 tunnel highlighted in Fig. 1a, with MED23 L343; another prominent edge joined MED23 L343 and MED15 L595. Thus, recurrent AD traffic was concentrated among a restricted set of receptor-site pairs on the Mediator Tail.

**Figure 3.**
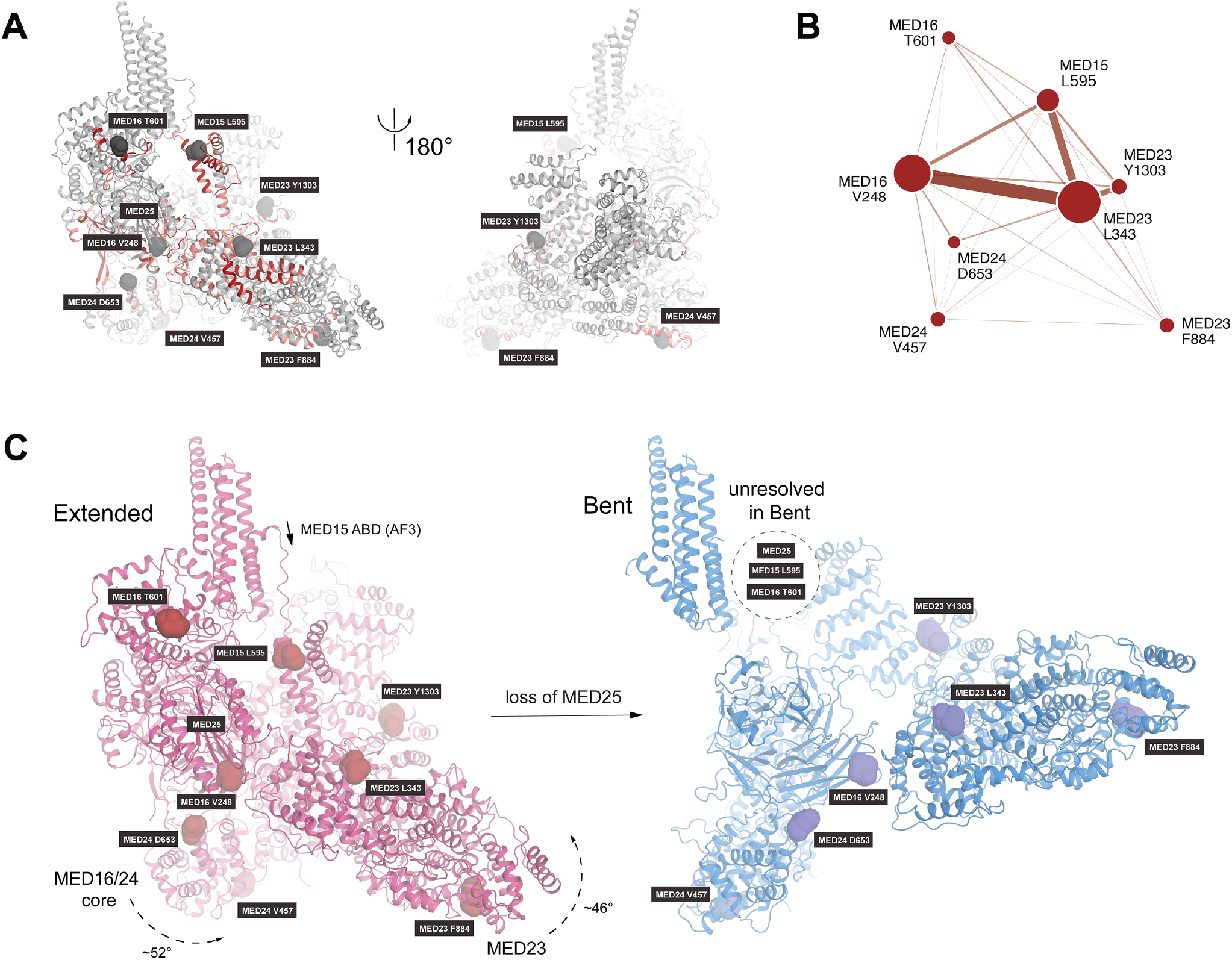
AD traffic converges on a Tail surface remodeled between Extended and Bent Mediator. **(A) AD traffic mapped onto the Extended Tail surface.** Two views of the Extended Tail, rotated by 180°. The Tail is shown in gray; recurrent AD-contact regions identified across the AF3 ensemble are highlighted in red, and labeled spheres mark recurrent AD-contact sites. **(B) Network of sites contacted by the same AD.** Node area reflects AD traffic at each site, and edge width the number of AF3 models in which the same AD contacts both connected sites under the strict hydrophobic criterion. The most frequent connection links MED16 V248 and MED23 L343, with another prominent edge between MED23 L343 and MED15 L595. **(C) Reorganization of AD-contact regions between Extended and Bent Mediator.** The Extended and Bent Tail structures are shown side by side in magenta and blue, respectively, after alignment through the shared MED15– MED27–MED29 helical bundle. In this reference frame, MED23 undergoes an approximately 46° rigid-body rotation, whereas MED24 undergoes an approximately 52° internal hinge-like reorientation. The MED15 ABD is shown at its AF3-modeled position in Extended Mediator. The dashed circle marks regions unresolved in Bent Mediator, including MED25, the MED16 region containing T601 and the MED15 ABD.

As noted above, the AF3 ensemble recovered only the Extended Tail form, so we next asked how these receptor sites are reorganized in the Bent form (Fig. 3c). After alignment through the shared Tail core, MED23 in the Bent Tail is rotated by approximately 46° relative to the Extended form, while MED24 undergoes an approximately 52° internal hinge-like reorientation. MED25 is likely dissociated in the Bent form [Chen et al., 2021], so the MED16–MED25 tunnel is not maintained; the MED16 helical region containing T601 is likewise unresolved. The MED15 ABD is unresolved experimentally in both forms and is shown at its AF3-modeled position in the Extended Tail (Fig. 3c); in the Bent form, accommodation of the unresolved MED16 helical domain is incompatible with retention of the MED15 ABD at its Extended-state position, suggesting that the ABD must also move upon the Extended-to-Bent transition. Thus, recurrent receptor regions are substantially reorganized in the Bent form through relative movement, displacement and loss of defined receptor architecture.

Importantly, this loss of structural definition in the Bent form extends beyond the Tail. In the global cryo-EM reconstruction, diffuse density is also evident in the distal Head–Middle region, whereas Extended Mediator is comparatively uniformly resolved [Chen et al., 2021] (Extended Data Fig. 3c). The focused Tail reconstructions provide a direct control showing that the lower structural definition of the Bent Tail is not due to fewer particle images contributing to the cryo-EM map. Local-mask classification and focused refinement yielded essentially matched particle populations for the Extended and Bent Tails—98,988 and 96,354 particles, respectively—yet the Extended Tail refined to 3.2 Å whereas the Bent Tail remained at 4.5 Å [Chen et al., 2021] (Extended Data Fig. 3a, b). This reduced structural definition is evident both in the Bent state of free Mediator and in PIC-bound Mediator reconstructions [Chen et al., 2021; Chen et al., 2022] (Extended Data Table 1). Persistent disorder strongly suggests that the Bent form is inherently more mobile throughout its length. This mobility extends from the TF-responsive Tail, where regulatory signals are received, to the Pol II-facing end, where Mediator acts on the initiation machinery.

Taken together, these comparisons reveal dramatic remodeling of the AD-engaging Tail surface upon the Extended-to-Bent transition. Large portions are repositioned, while others are displaced or lose definition in the Bent form. AF3 does not resolve state preference for individual ADs, but these structural differences suggest how multivalent AD binding could reshape the conformational landscape of the Mediator Tail.

### Serial helical paths suggest a route for long-range coupling through Mediator

Viewed in the context of the entire human Mediator complex, the Tail lies at one end of a series of elongated helical bundles extending toward Pol II-facing regions (Fig. 4a,b). At the Tail-proximal end, the first bundle incorporates helices from MED15 together with MED27 and MED29; successive bundles alternate in orientation as they course around MED14, then progressively diverge from it as they approach the Pol II clamp region. The resulting arrangement resembles an articulated mechanical linkage, with well-defined helical bundles joined by flexible loops, allowing relative motion to propagate through the assembly.

**Figure 4.**
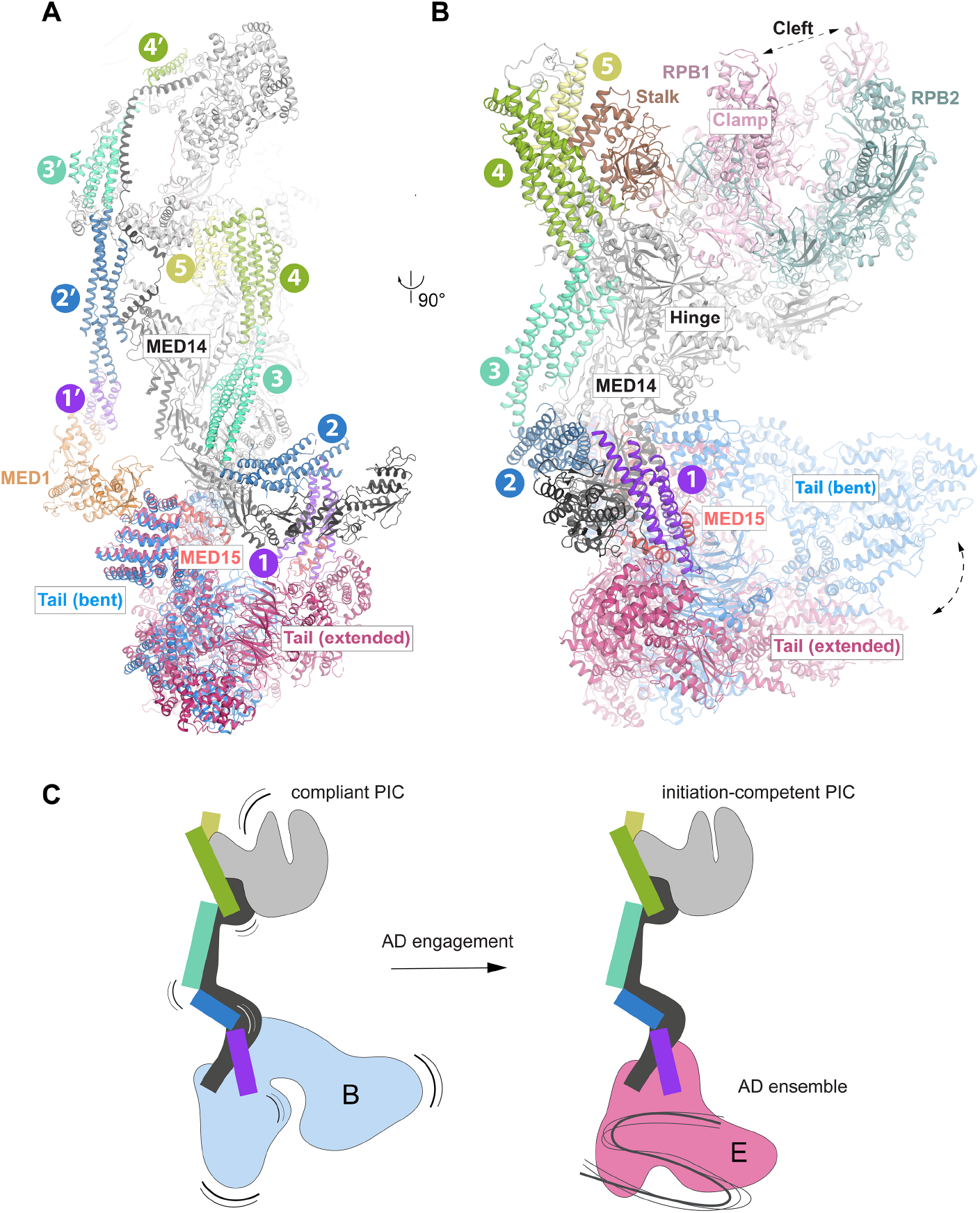
Serial helical paths suggest a route for coupling Tail state to PIC conformation. **(A) Helical paths spanning Mediator.** Back view of Mediator from the human +1 nucleosome–PIC–Mediator structure (PDB 8GXS) [Chen et al., 2022]. Pol II, the remaining PIC components and the nucleosome are omitted for clarity; the Bent Tail from PDB 7ENJ [Chen et al., 2021] is overlaid for comparison. Along the 1–5 path, bundle 1 comprises MED27/29 together with C-terminal MED15 helices; bundles 2–5 correspond approximately to MED27/29–MED28/30, MED28/30–MED11/22, MED11/22–MED8/17 and MED8/17, respectively. The 1′–4′ path begins near the MED1–Tail interface and traverses the Middle module toward the Mediator plank and a distinct Pol II-facing region, including the MED4–MED9 and MED7–MED21 bundles. **(B) The 1–5 path at the Mediator–Pol II interface.** Side view rotated by approximately 90° from panel A, with much of the remaining Mediator structure omitted for clarity. The path courses around MED14 to the stalk, beside the rear hinge of the Pol II clamp. RPB1, RPB2, the clamp and the cleft are indicated. Hinge marks the MED18– MED20/RPB3–RPB11 interface. **(C) Model for coupling Tail state to Pol II cleft geometry within the PIC.** Colored segments represent the 1–5 helical relay shown in panels A and B; the 1′–4′ path is omitted. MED14 and the Pol II-facing hinge are shown in black, and Pol II in gray, with the clamp on the left side of the cleft. Motion marks denote proposed broader conformational sampling at flexible bundle junctions, including greater motion of the Tail and Pol II clamp. In this model, AD engagement biases the Tail toward the more constrained Extended state, narrowing the range of configurations along the relay and favoring an initiation-competent PIC conformation, including the closed Pol II clamp associated with promoter melting and productive initiation.

At its Pol II-facing end, the helical path reaches the Mediator–Pol II interface at the stalk, beside the rear hinge of the Pol II clamp. This places the Tail and the clamp pivot region at opposite ends of the same articulated path, providing a direct structural route for coupling Tail state to the conformation of the initiation machinery.

A second, architecturally related helical path extends along the opposite side of MED14. Beginning near the MED1–Tail interface, it traverses the predominantly helical Middle module toward the Mediator plank and a distinct Pol II-facing region (Fig. 4a). Together, the two paths span Mediator from TF-responsive regions to distinct interfaces on the Pol II-facing core. Viewed alongside the state-dependent mobility described above, this architecture supports a model in which the Bent state samples a broader range of Pol II-facing configurations, while the more ordered Extended state constrains that range.

## Discussion

Structural and biochemical studies have highlighted two properties of Mediator that are central to transcriptional activation. The complex is structurally pliable, with substantial Tail mobility [Tsai et al., 2014; Zhang et al., 2021; Chen et al., 2021], and engages ADs through weak, dynamic and multivalent contacts [Brzovic et al., 2011; Tuttle et al., 2018; Sanborn et al., 2021]. The molecular basis connecting these two aspects has remained obscure. A mechanistic model of Mediator-dependent activation must therefore address two related questions: how the sequence and spacing of AD interaction elements are read by the Tail, and how the consequences of this engagement are propagated through Mediator to the Pol II initiation machinery. Near-atomic structures of mammalian Mediator and Mediator–PIC assemblies now provide a framework in which these questions can be considered [Zhao et al., 2021; Abdella et al., 2021; Chen et al., 2021; Rengachari et al., 2021].

We examined human AD–Tail interactions collectively through the lens of AlphaFold, providing a population-level view of the first question and a basis for considering the second. We used a large, functionally characterized set of human ADs as probes of the Tail surface and focused on features recurring across the ensemble, without assigning significance to individual poses. Our analysis revealed a recurrent interaction surface spanning multiple Tail subunits, frequently engaged by individual ADs at more than one site. Systematic deletion data broadly supported the structural view: regions required for activation often contacted the Tail in multiple models. By contrast, activation-enhancing deletions frequently shortened intervening linkers, arguing that activation depends on more than simply the sum of individual Tail contacts, with the spacing between Tail-engaging segments also contributing. Most importantly, the Tail surface sampled by ADs is reorganized between the Bent and Extended conformations [Chen et al., 2021], hinting at how ADs could program Tail motions.

In this view, individual ADs need not induce distinct activator-specific Mediator conformations, despite an earlier report [Taatjes et al., 2002]; their distributed contacts could instead reshape the Tail conformational landscape [Tsai and Nussinov, 2011]. A common theme across allosteric systems is that binding of a regulatory ligand redistributes a broad unliganded conformational ensemble toward a smaller set of functionally productive states [Boehr et al., 2009]. Mediator may exploit the same principle on a larger scale. In the Bent form, density is diffuse in the distal Tail and in the Pol II-facing Head–Middle region, whereas the Extended form is comparatively uniformly resolved (Extended Data Fig. 3c) [Chen et al., 2021; Kornberg and Feklistov, 2025]. For the Tail, particle number does not account for this difference: focused classification recovered essentially identical numbers of Extended and Bent Tail particles, yet the Bent Tail remained markedly less resolved, with several structural elements visible in the Extended Tail lacking defined density (Extended Data Fig. 3a,b). We therefore interpret the Bent state as a broader and more mechanically compliant ensemble associated with basal transcription, and the Extended state as a more constrained, initiation-competent form stabilized by activator engagement.

We consider it noteworthy that the Bent form has not been reported in earlier structures of endogenously expressed mammalian Mediator and emerged only in the recombinant, overexpressed human complex used by Chen et al. [El Khattabi et al., 2019; Zhao et al., 2021; Chen et al., 2021]. This may be explained by sample history. Strong TF–Mediator interactions persist through purification, as exemplified by the original isolation of human Mediator with thyroid hormone receptor and later recovery of the complex with other activators [Fondell et al., 1996; Kornberg, 2005 and references therein]. At the same time, bound activators remain invisible by cryo-EM, as illustrated by VP16 in the human Mediator–PIC complex [Abdella et al., 2021]. Retention of endogenous TFs may therefore have biased earlier preparations toward the Extended form, whereas overexpression could leave a greater fraction of Mediator free of activators, revealing more of the underlying Bent–Extended equilibrium.

The serial helical paths spanning Mediator form a plausible mechanical relay from the Tail to the Pol II-facing core [Tsai et al., 2017; Zhao et al., 2021]. The architecture itself is strikingly suggestive of long-range conformational coupling, through which changes in Tail compliance could reach Pol II. We favor cleft geometry as a key variable [Kornberg and Feklistov, 2025]. Mediator is positioned to influence both the clamp and the adjacent shelf module through the Head–stalk and Middle interfaces, respectively [Tsai et al., 2017; Kornberg and Feklistov, 2025]. Clamp opening was previously proposed from a complex with only the Mediator Head [Cai et al., 2012]. We instead favor the closed clamp, required for promoter melting and the formation of the first phosphodiester bond in multisubunit RNA polymerases [Feklistov et al., 2017; Dienemann et al., 2019; Chen et al., 2020; Mazumder et al., 2021; Feklistov, 2024], as the Pol II state promoted by the Extended form (Fig. 4c). Mediator–TFIIH interactions could be similarly modulated, affecting DNA entry into the cleft, contacts with the +1 nucleosome and RPB1 C-terminal domain phosphorylation required for promoter escape. [Chen et al., 2021; Chen et al., 2022; Abdella et al., 2021; Rengachari et al., 2021].

Bacterial transcription provides an evolutionary precedent for initiation control via reprogramming polymerase motions. The stress regulators TraR and DksA/ppGpp reshape the active-site cleft and restrict clamp motion toward the melting-competent closed state [Chen et al., 2019; A. Feklistov, unpublished observations], with promoter-dependent consequences for initiation kinetics. In bacteria, the clamp and shelf move together as part of a larger swivel module [Kang et al., 2018]. Because these modules are conserved across multisubunit RNA polymerases and are directly engaged by the helical paths, Mediator may have evolved to greatly elaborate this regulatory logic, decoding rich AD sequence grammar into probabilistic control of Pol II cleft geometry and productive initiation.

The distributed Tail-engagement model helps explain how combinations of DNA-bound activators may act cooperatively on a single Mediator complex [Zhou et al., 2025]. A central feature of multivalency is that individual contacts are interdependent: engagement at one site can influence engagement at others, with avidity and competition shaping their combined contribution within the same assembly [Holehouse and Kragelund, 2024; Sipko et al., 2024]. Multivalency may also account for the occurrence of bifunctional effector domains, in which the same short sequence supports both activation and repression [DelRosso et al., 2023], if alternative modes of engagement bias the Tail conformational landscape differently. The local ensemble of ADs could therefore collectively program Tail motions, allowing combinatorial TF input at an enhancer to tune promoter state. This model does not exclude recruitment of Mediator to the promoter by ADs as a contributor to PIC assembly.

Our model makes several testable predictions, although directly resolving transient AD contacts within a megadalton complex and parsing their effects on its conformational landscape will likely remain a formidable challenge even as structural methods improve. The present analysis delineates relationships between AD architecture, Tail state and Pol II geometry that provide concrete experimental entry points for future mechanistic studies of eukaryotic transcription regulation.

## Extended Data

**Extended Data Fig. 1.**
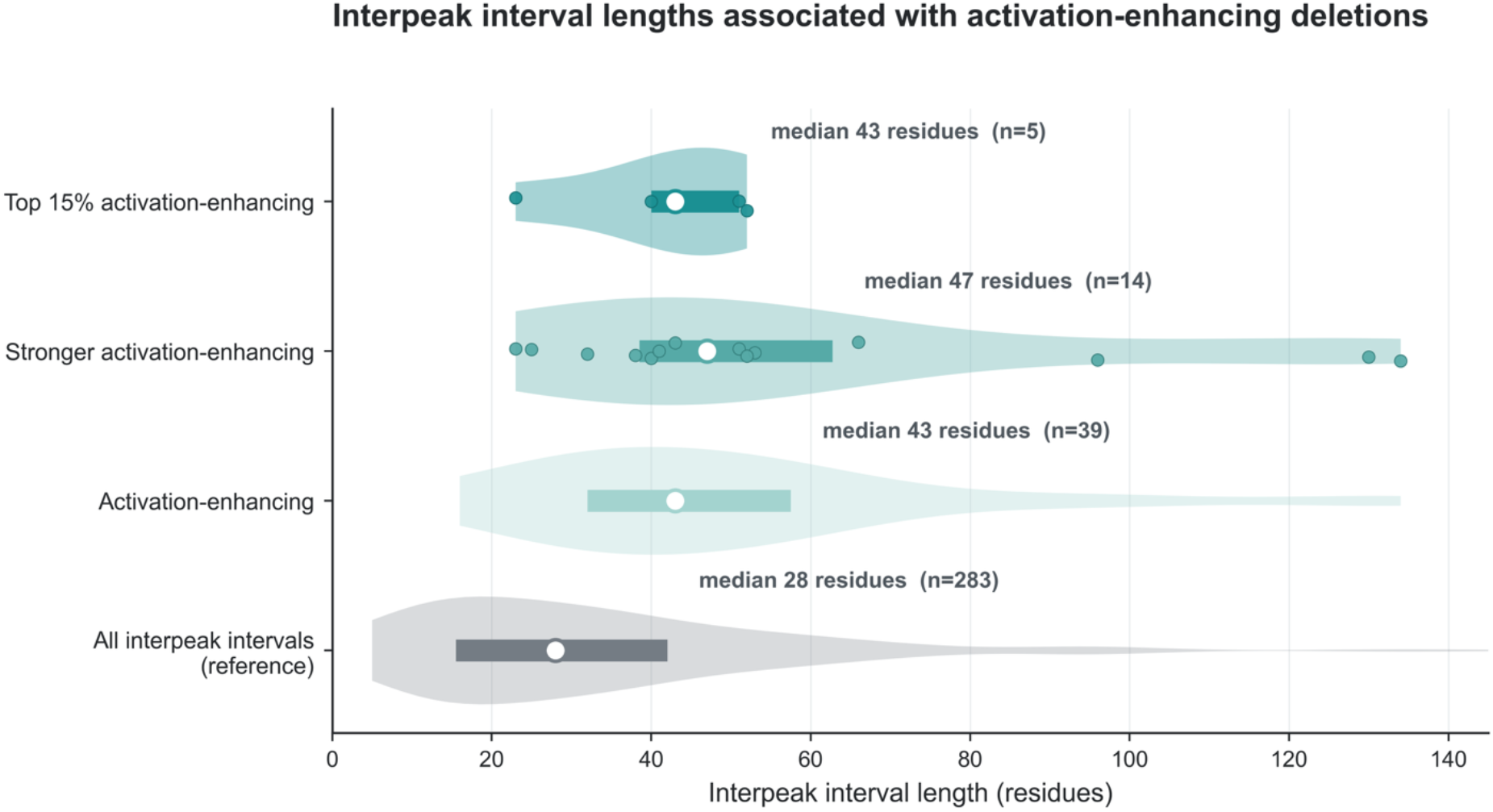
Activation-enhancing deletions are associated with longer interpeak intervals. Distribution of sequence distances between adjacent peaks of Mediator contact, shown for all peak pairs (reference; n = 283) and for those associated with activation-enhancing deletions (n = 39), including stronger (n = 14) and top 15% (n = 5) subsets. Violin plots show the interval-length distributions; thick horizontal bars indicate interquartile ranges, open circles indicate medians, and filled points show individual intervals only for groups containing ≤15 intervals. Median interval lengths were 28 residues for all interpeak intervals, 43 residues for intervals associated with activation-enhancing deletions, 47 residues for the stronger subset, and 43 residues for the top 15% subset. Some modeled constructs directly concatenate sequence segments separated in the native protein; intervals crossing these joins therefore do not represent native sequence spacing. Phenotype thresholds and the definition of interpeak intervals are described in Methods.

**Extended Data Fig. 2.**
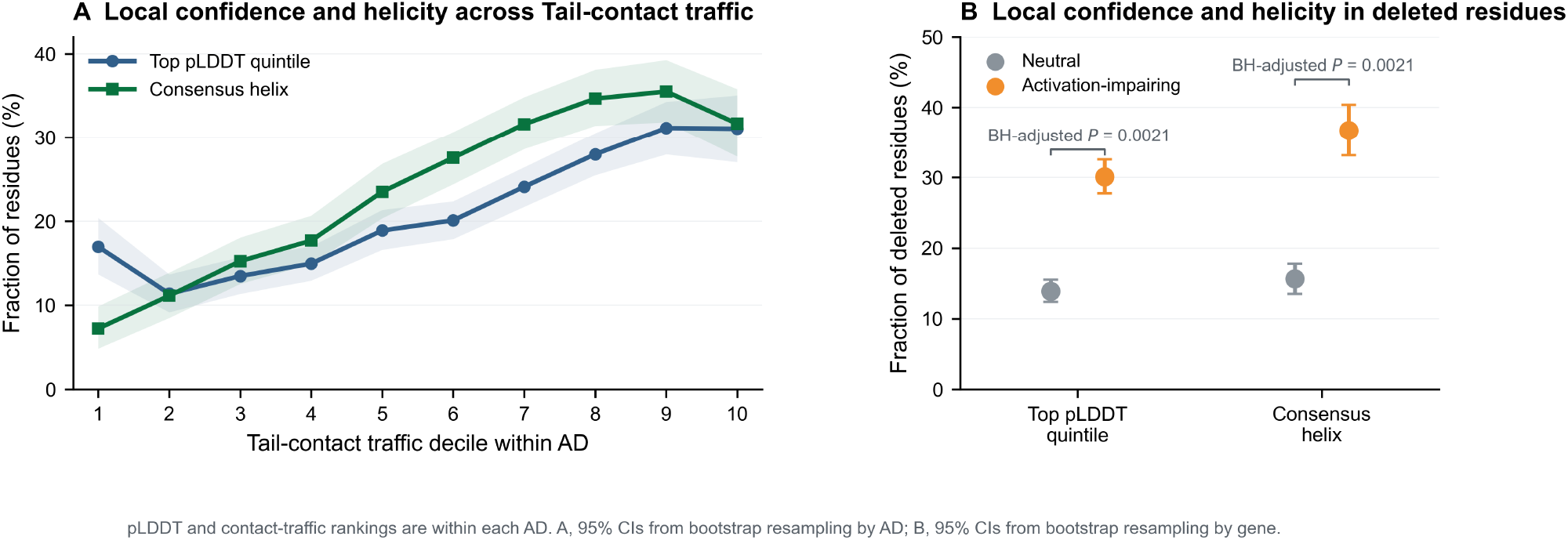
Local AF3 confidence and helicity are enriched in Tail-contacting AD sequence and regions removed by activation-impairing deletions. **(A) Local confidence and helicity across Tail-contact traffic.** Residues were ranked by Tail-contact traffic within each AD and divided into deciles. Points show the fraction of residues in each decile that fall within the highest 20% of pLDDT values in that AD (blue) or within a consensus helix (green), with equal weighting across ADs. The analysis included 25,650 residues across 234 modeled ADs. Consensus helices were defined as residues called helical by STRIDE in at least 40% of usable models and occurring in runs of at least three residues. **(B) Local confidence and helicity in regions removed by deletions.** Points show the fraction of residues removed by neutral deletions (1,825 windows from 219 genes) or activation-impairing deletions (1,639 windows from 208 genes) that fall within the highest 20% of pLDDT values in the corresponding AD or within a consensus helix. The activation-impairing group includes deletions that abolished activation. Deletion windows without corresponding pLDDT values were excluded. Shading (A) and error bars (B) indicate 95% confidence intervals. Both comparisons yielded Benjamini–Hochberg-adjusted *P* = 0.0021. AF3 pLDDT reports confidence in local AD conformation, not in its placement on Mediator.

**Extended Data Fig. 3.**
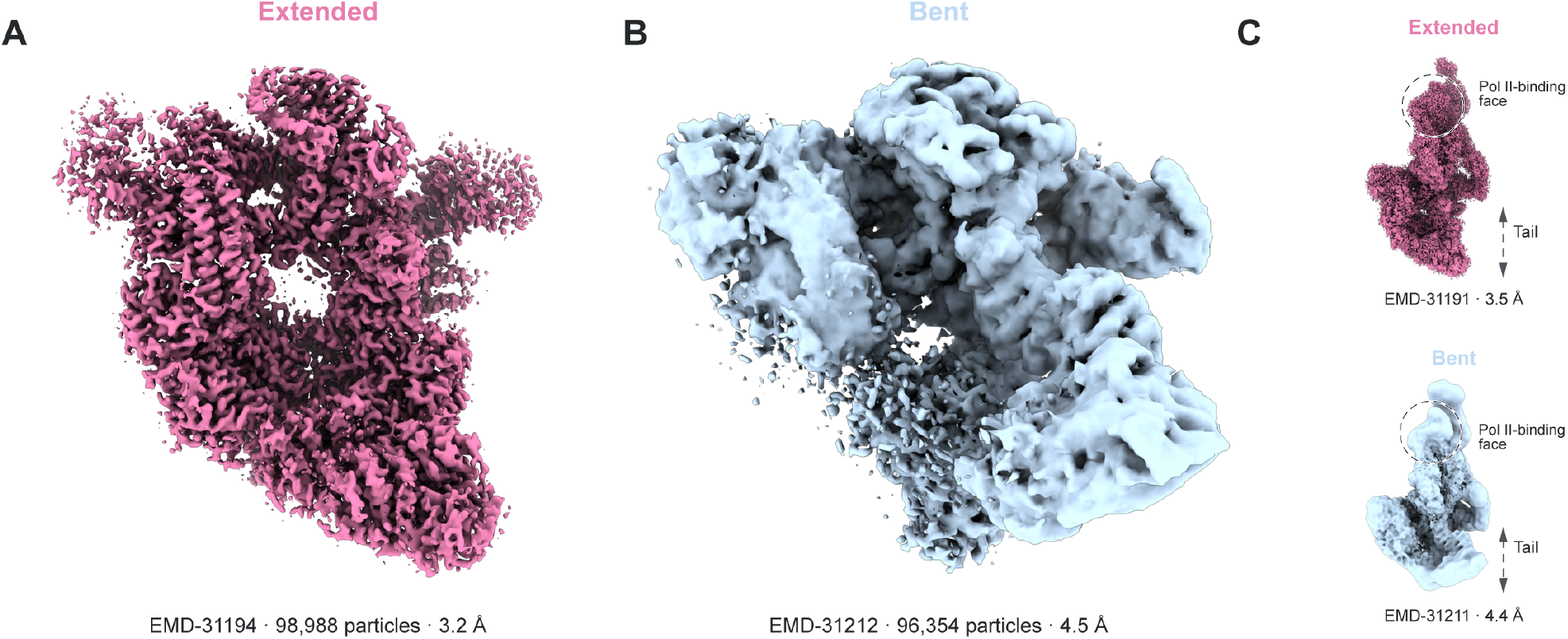
Reduced structural definition of Tail-bent Mediator in cryo-EM reconstructions. **(A,B) Focused Tail-module reconstructions** of the Extended (MEDᴱ; EMD-31194, 3.2 Å; 98,988 particles) and Bent (MEDᴮ; EMD-31212, 4.5 Å; 96,354 particles) states of ΔMED1-IDR Mediator. The lower resolution of MEDᴮ despite nearly matched particle populations is consistent with greater conformational heterogeneity of the Bent Tail. **(C) Overall reconstructions** from the same ΔMED1-IDR Mediator dataset: MEDᴱ (EMD-31191, 3.5 Å) and MEDᴮ (EMD-31211, 4.4 Å). Compared with MEDᴱ, the MEDᴮ reconstruction shows more diffuse density in the Tail and Pol II-binding face, consistent with greater conformational heterogeneity at both ends of the complex. Cryo-EM maps are from Chen et al., 2021.

**Extended Data Table 1.**
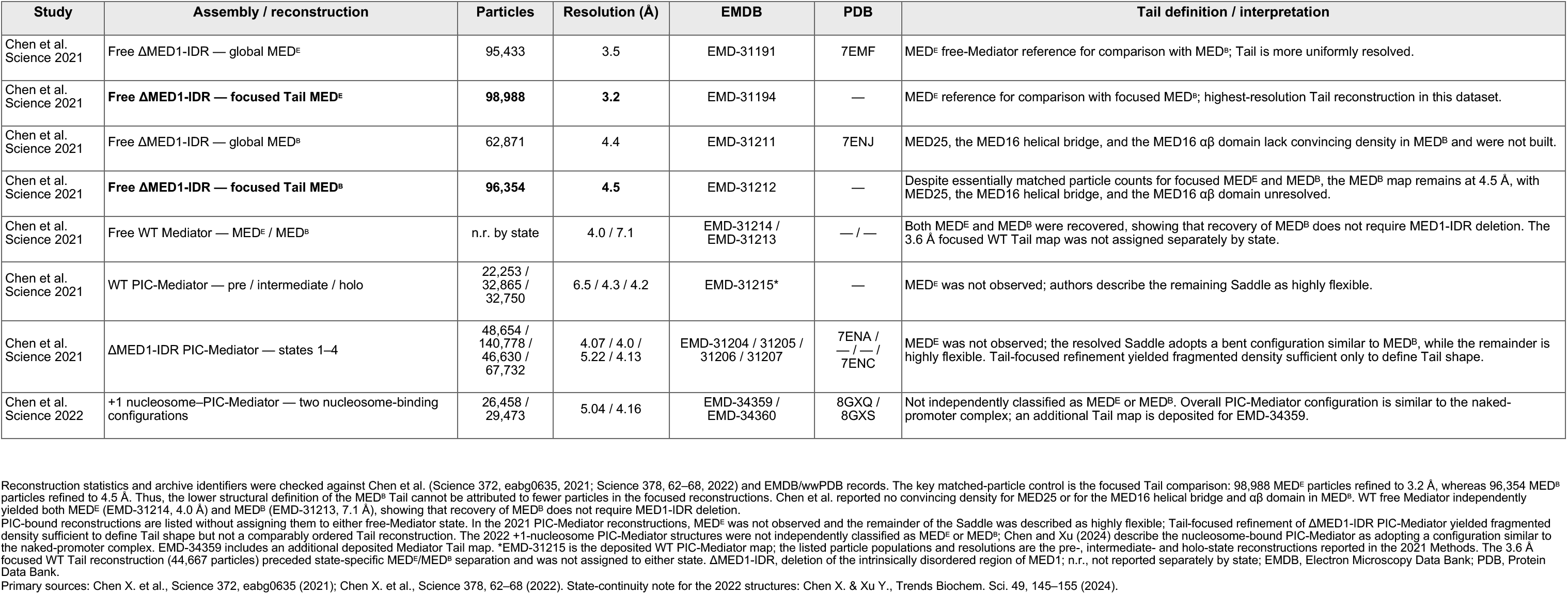
Cryo-EM reconstruction statistics and Tail structural definition across free and PIC-bound human Mediator reconstructions.

## Materials and Methods

### Activation-domain dataset and modeled constructs

Human activation-domain (AD) annotations and functional measurements were obtained from DelRosso et al. [2023]. In that study, protein sequences were screened as overlapping 80-aa tiles with a 10-aa step, and ADs were defined from contiguous activating tiles. We used all 374 AD intervals reported in Supplementary Table 2 (“Activation Domains”), representing 293 unique HGNC symbols. For each gene, all reported AD intervals were ordered by native protein coordinate and combined into a single modeling construct. Overlapping residues were included once; adjacent or separated intervals were concatenated directly, with native residues between separated intervals omitted. This yielded 293 modeled constructs, including 22 assembled from two or more noncontiguous native regions. The merged sequences were used as modeling inputs and were not intended to redefine biological AD boundaries.

### AlphaFold 3 modeling of AD–Mediator Tail complexes

AD–Mediator Tail complexes were modeled with AlphaFold 3 (AF3) [Abramson et al., 2024] using AlphaFold Server. The fixed seven-chain Mediator receptor comprised MED15 (Q96RN5, residues 520–653), MED27 (Q6P2C8, residues 7–77), MED29 (Q9NX70, residues 55–136), MED16 isoform 2 (Q9Y2X0-2, residues 1– 841), MED25 (Q71SY5, residues 14–231), MED24 (F5GY88, residues 1–1008) and MED23 (Q9ULK4, residues 1–1368). The MED15 fragment retained the Tail-proximal structured activator-binding domain. Each AD was modeled as an additional chain. The 293 jobs produced five models each, yielding 1,465 predicted complexes. All models were carried forward to the artifact-filtering procedure described below.

### Artifact filtering for ensemble analysis

Models were screened for recurrent AF3 artifacts in which AD sequence was incorporated into missing Mediator structure (structural inpainting) or passed through sterically implausible regions of the Tail assembly (threading). An initial scripted inpainting screen identified severe AD penetration after receptor alignment. Models with at least 10 AD heavy atoms within 1.2 Å of the receptor blocker set were excluded when the alignment RMSD was ≤8 Å; models with less certain alignments were flagged for review. This screen excluded 86 of the 1,465 models.

A subsequent artifact review combined visual inspection with scripted application of a curated artifact ledger. Models dominated by inpainting were excluded from primary interface and contact analyses, whereas localized inpainting was handled by masking only the affected AD residues. Candidate threading regions were identified from heavy-atom overlap with the remainder of the complex and confirmed by visual inspection.

Confirmed threading regions were masked locally, with the remainder of each model retained. Applied to the remaining 1,379 models, this review excluded 240 additional models from interface and contact analyses and masked 913 model-specific AD-residue instances affected by partial inpainting and 61 affected by local threading. The resulting dataset comprised 1,139 models from 234 AD constructs.

AF3 ipTM/ipSAE interface-confidence metrics were evaluated separately as technical quality controls and were not used for model selection. Because pairwise interface-confidence scores were generally low, AF3 outputs were interpreted at the ensemble level as recurrent surface-usage tendencies rather than as uniquely resolved docked poses.

### Contact analysis and receptor-site definition

AD–Tail contacts were measured separately in each retained AF3 model. AD residues affected by the local inpainting or threading artifacts described above were omitted from these calculations. Proximity contacts were defined by Cα–Cα distances <12 Å. Strict hydrophobic contacts required both the AD and Tail residue to be W, F, Y, L, I, V or M, with at least one pair of non-hydrogen side-chain atoms separated by ≤4.5 Å.

To map recurrent AD contact across the Tail, proximity contacts were weighted continuously by distance using (1 − (d/12)²)², where d is the Cα–Cα distance in Å, so that close contacts contributed more strongly than contacts near the 12-Å cutoff. For each AD construct, scores were summed across retained models and AD residues for each Tail residue and normalized by the total contact score for that construct. The normalized maps were then averaged across constructs, giving each construct equal weight in the aggregate Tail-contact map. The same distance weighting was retained for strict hydrophobic contacts.

Recurrent receptor sites were defined from the aggregate Tail-contact map. Contacted residues were grouped separately on each Tail subunit when they were close in sequence, within three residues, or when their contact-weighted mean Cα positions were within 10 Å. Candidate site centers comprised residues at or above the 80th percentile within a candidate region or the 90th percentile globally; additional residues at or above the 50th percentile within the region or the 70th percentile globally were included in the candidate site envelope.

Candidate sites were then reviewed in structural context to merge related regions and refine site boundaries. Final receptor-site definitions are provided in the Source Data and were used for site-level traffic and co-engagement analyses.

### Analysis of multisubunit AD–Tail contacts and receptor-site co-engagement

For each usable AF3 model, retained post-mask AD–Tail contacts were used to score Tail subunits independently under the proximity and strict hydrophobic criteria defined above. A Tail subunit was counted as contacted when it contained at least one qualifying AD–Tail residue pair. An AD construct was classified as showing recurrent multisubunit engagement when at least two of its usable models each contacted two or more Tail subunits; the identities of the contacted subunits were not required to be the same across models.

Receptor-site co-engagement was evaluated within individual usable models using the strict hydrophobic criterion. A receptor-site pair was scored as co-engaged when the AD made at least one retained strict hydrophobic contact with each site envelope. In the receptor-site network, node area represents total retained strict hydrophobic contact mass within each envelope, and edge width represents the number of usable models co-engaging each site pair.

### Analysis of functional deletion phenotypes and AD–Mediator contact profiles

Systematic deletion measurements from DelRosso et al. [2023] were mapped from native protein coordinates onto the corresponding retained sequence segments of each modeled AD construct; no analyzed deletion window crossed a concatenation junction. Among deletion windows with complete measurements and stable expression (|ΔFLAG| ≤ 1.0), activation-impairing windows had ΔminCMV ≤ −1.0, activation-enhancing windows had ΔminCMV ≥ 1.1206, and neutral windows lay between these thresholds. A stronger activation-enhancing subset had ΔminCMV ≥ 1.6206, and the top 15% of activation-enhancing windows had ΔminCMV ≥ 2.2602. Deletions that abolished activation in DelRosso et al. were retained as a separate annotation. For Fig. 2c, the activation-impairing group comprised the threshold-defined activation-impairing windows together with deletions that abolished activation; Fig. 2d used only the threshold-defined class.

Residue-level Mediator contact mass was summed across Tail subunits and usable models, with residues lacking retained contacts assigned zero. The resulting profile was smoothed with a centered seven-residue mean and converted to an average-rank percentile within each AD. Deletion-centered profiles spanning −30 to +30 residues around the rounded deletion-window center were averaged by class and smoothed with a centered five-position mean for display.

Contact peaks were defined as contiguous runs of at least two residues exceeding the larger of the AD-specific 85th percentile of positive seven-residue-smoothed contact mass or 25% of that AD’s maximum smoothed value. Interpeak intervals were gaps of at least five residues between adjacent peaks. A deletion was associated with an interval when it overlapped the interval by at least five residues or by at least 50% of the deletion length; each interval was counted once per deletion class. For the 22 constructs containing noncontiguous native sequence, interpeak distances were measured in modeled-construct coordinates, and intervals spanning concatenation junctions therefore do not represent native sequence spacing.

For Fig. 2d, a deletion window was classified as WFYL-free when it contained none of W, F, Y or L. Confidence intervals were the 2.5th and 97.5th percentiles of 5,000 gene-level bootstrap resamples, with all deletion windows from each sampled gene retained together.

### Local AF3 confidence and secondary-structure analysis

For each AD, residues were ranked by pLDDT within the modeled construct, and the highest quintile was designated the top-pLDDT group. The analysis included 25,650 residues across 234 modeled ADs.

Secondary structure was assigned independently in each usable AF3 model using STRIDE [Frishman and Argos, 1995]. Residues assigned to the STRIDE α-helical state (H) in at least 40% of usable models were classified as consensus helical when they occurred in contiguous runs of at least three residues.

For Extended Data Fig. 2a, residues were grouped by Tail-contact traffic decile within each AD. The fractions of residues belonging to the top pLDDT quintile or to consensus helices were calculated within each AD and averaged with equal weighting across ADs. Confidence intervals were obtained from 5,000 bootstrap resamples by AD.

For Extended Data Fig. 2b, only deletion windows with mapped pLDDT values were included. The fractions of deleted residues belonging to the top pLDDT quintile or to consensus helices were calculated for neutral and activation-impairing deletions. Confidence intervals and two-sided P values were obtained from 5,000 gene-level bootstrap resamples, retaining all deletion windows from each sampled gene. P values were adjusted by the Benjamini–Hochberg procedure across the combined family of 36 deletion-class/metric comparisons.

### Comparison of Extended and Bent Tail conformations

The Extended and Bent human Mediator structures (PDB 7EMF and 7ENJ, respectively) from Chen et al. [2021] were superposed using corresponding Cα atoms from the shared MED15–MED27–MED29 helical bundle.

MED23 rotation was measured by least-squares superposition of 1,334 sequence-matched Cα atoms from the two structures, yielding a rotation of 46.13°. MED24 hinge motion was analyzed by treating MED24 as two rigid segments separated by a single hinge. Candidate hinge positions were tested along the matched sequence, with at least 60 Cα pairs required in each segment, and the position giving the lowest combined fitting residual was selected. The relative rotation between the two segments was 52.14°.

MED25 and the MED16 region containing the T601-centered receptor site are unresolved in the Bent structure and were therefore not assigned Bent-state receptor positions.

### Comparison of predicted Tail geometry with an experimental structure

Model 0 from each of the 293 AF3 prediction sets was compared with the experimental human Mediator structure PDB 8GXS [Chen et al., 2022]. Residue correspondence for MED15, MED29, MED16, MED25, MED24 and MED23 was established by sequence alignment. All 3,328 matched Cα atoms were included in a single least-squares superposition without outlier rejection. Across the 293 comparisons, the median Cα RMSD was 1.892 Å (interquartile range, 1.786–1.963 Å).

### Generative AI use

Generative AI tools (OpenAI ChatGPT and Codex; Anthropic Claude Code) were used in the development of custom analysis scripts, repository and provenance checks, review of analysis logic and outputs, and manuscript editing. Scientific interpretation and evaluation of the reported results remained the responsibility of the authors.

### Data availability

Human activation-domain sequences and functional measurements were obtained from DelRosso et al. [2023]. Experimental structural coordinates are available from the Protein Data Bank under accession codes 7EMF, 7ENJ and 8GXS. Processed data underlying the quantitative analyses, receptor-site definitions and artifact-filtering assignments are available in the Zenodo archive: https://doi.org/10.5281/zenodo.23088675.

### Code availability

Custom scripts used for the reported analyses are available at https://github.com/andreyf2022/mediator-ad-tail-analysis. The version used for this study is available as GitHub release v1.0.1 and archived at Zenodo: https://doi.org/10.5281/zenodo.23088675.

## Acknowledgements

A.F. is grateful to Lacramioara Bintu and Nicole DelRosso for helpful discussions and insights into the activation-domain dataset.

## Author Contributions

A.F. conceived the mechanistic hypothesis underlying the study and developed the conceptual framework. A.F. performed the analyses and prepared the manuscript and figures. R.D.K. contributed to scientific discussions during development of the work and commented on the manuscript. Both authors reviewed and approved the manuscript.

